# Distinct mechanisms decommission redundant enhancers to facilitate phenotypic evolution

**DOI:** 10.1101/2025.10.27.684981

**Authors:** Areej Said-Ahmad, Noa Shimron, Ela Fainitsky, Sujay Naik, Srijani Roy, Nicolás Frankel, Ella Preger-Ben Noon

**Affiliations:** Department of Genetics and Developmental Biology, The Rappaport Faculty of Medicine and Research Institute, Technion-Israel Institute of Technology, Haifa, Israel; Instituto de Fisiología, Biología Molecular y Neurociencias (IFIBYNE), Consejo Nacional de Investigaciones Científicas y Técnicas (CONICET) y Universidad de Buenos Aires (UBA), Buenos Aires 1428, Argentina

**Author notes:** **Corresponding author:** Ella Preger-Ben Noon.

## Abstract

The evolutionary loss of morphological traits is often driven by changes in gene regulation. Many developmental genes are controlled by multiple, redundant enhancers, raising the question of how robust regulatory systems can be dismantled to permit phenotypic transitions. We show that the loss of larval trichomes in *Drosophila sechellia* resulted from the independent inactivation of four embryonic enhancers of the *shavenbaby* gene. Each enhancer was extinguished by a distinct mechanism: (1) a large deletion that removed essential sequences, (2) the loss of activator sites and gain of repressor sites, (3) the acquisition of a long-range silencer, and (4) the unmasking of pre-existing repression. Notably, three of these mechanisms relied on repression, pointing to repression as a rapid route for the evolutionary loss of robust regulatory elements. These results show that robustness in gene regulation does not prevent morphological change but instead provides multiple opportunities for mutations to reduce enhancer activity, giving selection many paths to reshape form.

## Introduction

The loss of morphological features has occurred frequently throughout the evolutionary history of life. This process, known as regressive evolution, usually occurs when certain traits become disadvantageous or neutral in a given environment (*1, 2*). Some of the best-known examples of adaptive regressive evolution include the loss of tails in primates (*3*), limbs in snakes (*4, 5*), eyes and pigmentation in cavefish (*1, 6*–*8*), pelvic spines in freshwater sticklebacks (*9, 10*), and spots in *Drosophila* wings (*11, 12*). Most of these evolutionary transitions involved sequence changes in *cis*-regulatory regions of key developmental genes, leading to reduction or loss of gene expression in a specific domain during development. For example, the repeated loss of the pelvic girdle in freshwater populations of sticklebacks resulted from deletions of a pelvic enhancer of the *Pitx1* gene (*9, 10*). Similarly, hindlimb loss in snakes evolved through the accumulation of single-base mutations and small deletions in the ZRS enhancer of *Sonic hedgehog* (*Shh*), which affected transcription factor binding sites (*4, 5*). Loss of wing spots in *Drosophila*, on the other hand, partially evolved through the acquisition of a silencer element near the spot enhancer of *yellow* (*11*). In all these cases, the loss of morphological features resulted from a loss of function of a single developmental enhancer. This mechanism is thought to be facilitated by the modular nature of *cis*-regulatory regions (*13*), allowing a single enhancer of a given gene to evolve without affecting the function of other enhancers, thereby minimizing pleiotropic effects (*14, 15*).

However, the modularity paradigm has been challenged by recent experimental data (reviewed in Kittelmann et al. 2021 (*16*) and Sabaris et al. 2019 (*17*)). For example, many developmental genes have multiple enhancers with redundant functions (*18*–*22*). These so-called “shadow enhancers” have been demonstrated to ensure robust gene expression in the face of genetic and environmental variability (*19*–*21*). Another level of robustness may exist at the level of individual enhancers that contain multiple binding sites for the same transcriptional activator (*23*–*26*). How this inherent robustness can be overcome to evolve new phenotypes remains unclear, as there are almost no examples of the loss of multiple redundant enhancers underlying an evolutionary change.

One exception to this is the partial loss of function of the *shavenbaby* (*svb*) gene in *Drosophila sechellia*. The *svb* gene encodes a transcription factor that controls the formation of non-sensory, hair-like structures, named trichomes, on the cuticle of insects. Work in *D. melanogaster* revealed that the embryonic expression of *svb* is controlled by seven enhancers (designated *7, E6, E3, A, Z1*.*3, DG3* and *DG2*) that are scattered across a 90 kilobase (kb) region upstream of the core promoter (*20, 27*–*29*) (Fig. 1). These enhancers drive both unique and overlapping expression patterns, providing robustness to *svb* expression under environmental or genetic variation (*20*). In addition, it has been shown that some of these enhancers contain multiple binding sites for the same transcription factor (*23, 24*).

**Fig. 1:**
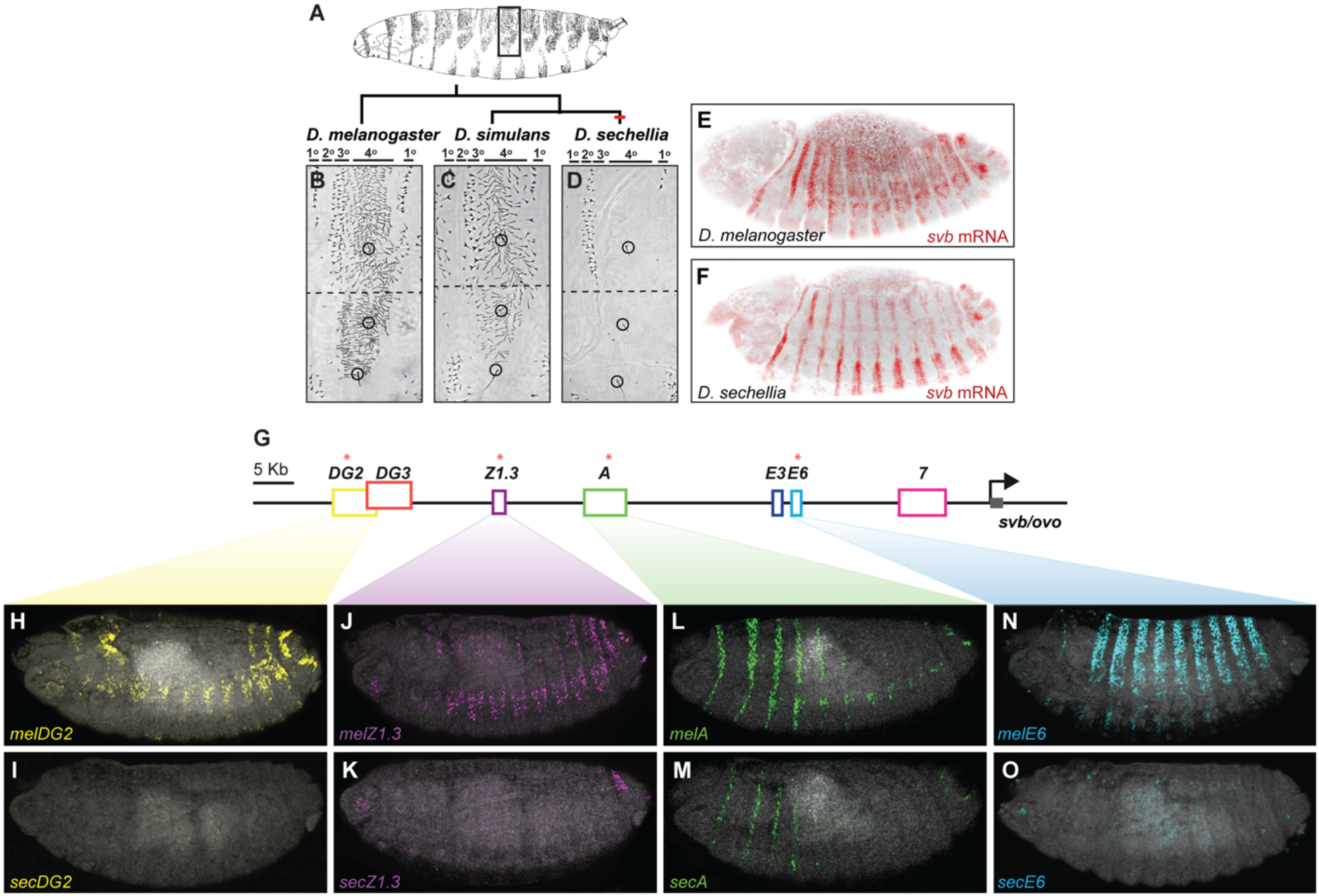
Trichome patterns have evolved between *Drosophila* species by changes in the regulatory region of the *shavenbaby* gene. **(A)** Lateral-view drawing of first instar larva of *D. melanogaster*. The rectangle marks the region shown in B-D. **(B-D)** Dorso-lateral cuticle of the fifth abdominal segment *D. melanogaster* (B), *D. simulans* (C), and *D. sechellia* (D). The circles indicate three sensory bristles, which are found in all three species. Adapted from McGregor et., 2007 (29). **(E-F)** Expression patterns of *svb* mRNA in stage 14 embryos of *D. melanogaster* (E) and *D. sechellia* (F) visualized by HCR™ RNA fluorescent *in situ* hybridization. **(G)** Schematic representation of the *svb* locus, indicating embryonic enhancers (colored boxes). The *svb* gene is marked with a gray arrow. The four enhancers that lost function in *D*.*sechellia* are marked with * **(H-O)** Expression of *D. melanogaster DG2::LacZ* (H, *melDG2*), *D. sechellia DG2::LacZ* (I, *secDG2*), *D. melanogaster Z1*.*3::LacZ* (J, *melZ1*.*3*), *D. sechellia Z1*.*3::LacZ* (K, *secZ1*.*3*), *D. melanogaster A::LacZ* (L, *melA*), *D. sechellia A::LacZ* (M, *secA*), *D. melanogaster E6::LacZ* (N, *melE6*), and *D. sechellia E6::LacZ* (O, *secE6*) reporter constructs in *D. melanogaster* stage 15 embryos.

Despite the intrinsic robustness of the *svb* regulatory region (*20, 23, 24, 30*), trichome patterns have repeatedly evolved in larvae of the genus *Drosophila* through *cis*-changes in *svb* regulation during embryogenesis (*31*–*35*). For example, the dorsal and lateral cuticle of first instar larvae of *D. melanogaster* and *D. simulans* exhibits broad patches of fine trichomes known as quaternary trichomes (Fig. 1A-C). In contrast, their sister species *D. sechellia* has lost these trichomes thus possessing a naked quaternary region (Fig. 1D). This morphological change is associated with reduced embryonic activity of four *svb* enhancers (Fig. 1H-O), leading to a loss of *svb* expression in quaternary cells of the embryonic epidermis (Fig 1 E-F) (*20, 27, 29*). We previously identified the genetic and molecular basis for the inactivation of the *svb E6* enhancer in *D. sechellia* (Fig. 1N-O) (*23, 36*). In *D. melanogaster, E6* encodes multiple binding sites for the transcriptional activators Arrowhead and Pannier, which are required for robust *svb* expression. In *D. sechellia*, nucleotide substitutions in the *E6* enhancer disrupted four Arrowhead binding sites and created a novel binding site for the transcriptional repressor Abrupt, resulting in complete loss of enhancer activity (*23*). Although we understand the inactivation of *E6* in detail, the genetic mechanisms underlying the loss of function of the other three enhancers remain unknown. This leaves an open question as to whether evolution operated through similar or distinct mechanisms to decommission different enhancers in a complex regulatory region.

Here, we uncover the genetic basis for the loss of activity of three additional *svb* enhancers in *D. sechellia*, providing the first complete portrait of the genetic events underlying a case of regressive evolution. Remarkably, each of the three *svb* enhancers lost activity through a different genetic mechanism: the *Z1*.*3* enhancer acquired a 120-nucleotide deletion in its core sequence, the *A* enhancer evolved through the acquisition of a silencer element, and the *DG2* enhancer lost an activator binding site, rendering it susceptible to pre-existing repression. These findings demonstrate that independent elements within robust and redundant regulatory regions can evolve through different genetic routes to converge on the same functional outcome.

## Results

### One large-effect deletion underlies the loss of the *Z1*.*3* enhancer activity in *D. sechellia*

The embryonic expression of *svb* in *D. melanogaster* is driven by seven enhancers, four of which have lost activity in *D. sechellia* (Fig. 1H-O). For example, the *D. melanogaster Z1*.*3* enhancer (*melZ1*.*3*), a 1.3 kb sequence located 61 kb upstream of the *svb* promoter, drives patches of expression in the dorsal and lateral epidermis of stage 15 embryos (Fig. 1J). In contrast, the orthologous sequence from *D. sechellia* (*secZ1*.*3*) drives expression only in the most posterior abdominal segment (Fig. 1K).

To identify the genetic changes responsible for the reduced activity of *secZ1*.*3*, we first performed a multiple sequence alignment, comparing the *secZ1*.*3* sequence to its orthologs from four related species that produce quaternary trichomes (Fig. S1). This analysis revealed sixteen single-nucleotide substitutions and deletions of 120 nucleotides and 35 nucleotides, specific to the *secZ1*.*3* enhancer (Fig. 2A and Fig. S1).

**Fig. 2:**
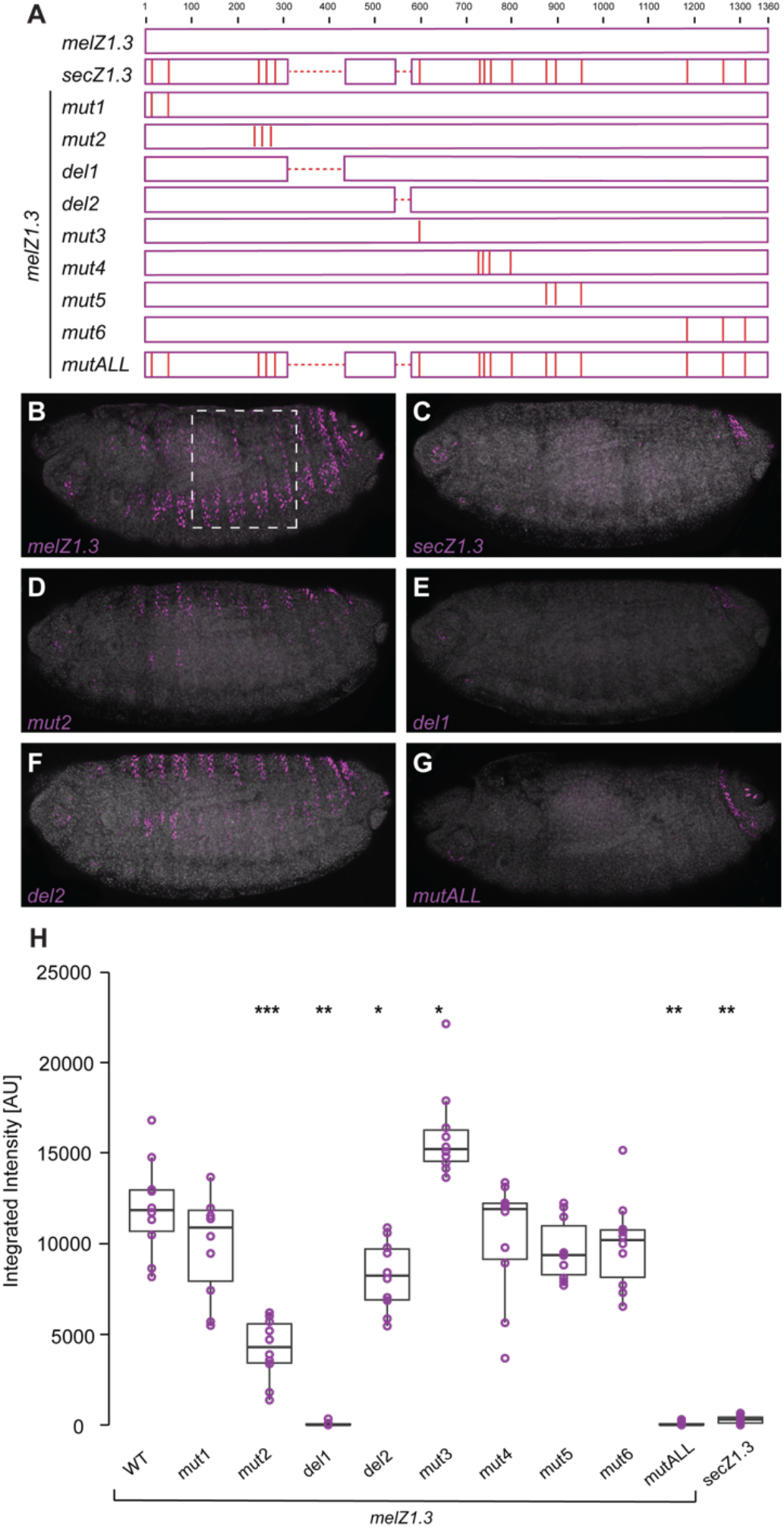
One large effect deletion and three small effect nucleotide substitutions underly the loss of *Z1*.*3* in *D. sechellia*. **(A)** Schematic representation of the mutated *Z1*.*3* enhancer reporter constructs. The positions of *D. sechellia*-specific substitutions and deletions are indicated with vertical and dashed red lines, respectively. **(B-G)** Expression of *melZ1*.*3* (B) and *secZ1*.*3* (C) reporter constructs and of *melZ1*.*3* reporter constructs carrying the indicated *D. sechellia*-specific deletions and substitutions in stage 15 embryos. **(H)** Box plots showing the integrated intensity of reporter activity in nuclei carrying the indicated constructs in the region outlined in B (n=10 embryos for each genotype). In the plots, each point represents an individual embryo. Asterisks denote significant difference from *melZ1*.*3* wild-type activities, * – P<0.05, **– P<0.01, ***– P<0.001 (Kruskal–Wallis test with Hochberg-adjusted post hoc comparisons).

To test the effect of these changes on enhancer function, we first introduced all *D. sechellia*-specific changes into the *melZ1*.*3* sequence and assessed its activity using reporter gene assays (Fig. 2). In stage 15 embryos, the resulting construct (*melZ1*.*3_mutALL*) drove little to no expression, similar to the *secZ1*.*3* enhancer (compare Fig. 2C and G). Next, we grouped the *D. sechellia*-specific substitutions into six clusters and introduced each cluster of mutations separately into *melZ1*.*3* (Fig. 2, Fig. S1 and Fig. S2). We also removed the nucleotides in *melZ1*.*3* that correspond to the two *D. sechellia*-specific deletions. This analysis revealed that the 120-nucleotide deletion is the primary cause of the reduced embryonic activity of *secZ1*.*3* (Fig. 2E and H). This deletion removes sequences from the core *Z1*.*3* enhancer, known as *Z0*.*3* (*28*). In addition, three nucleotide substitutions, also within this core element (*melZ1*.*3_mut2*; Fig. 2D andH), caused a slight but significant reduction in *melZ1*.*3* activity.

Together, these results indicate that the loss of embryonic activity of *Z1*.*3* in *D. sechellia* most likely resulted from a large-effect deletion within the core enhancer, which removed essential activator binding sites. It is not possible to disentangle whether the substitutions that affect the activity of the enhancer occurred before or after the deletion. In any case, this mechanism is markedly different from that of the *E6* enhancer, which evolved reduced function in the *D. sechellia* lineage through single-nucleotide substitutions that disrupted activator binding sites and created a site for a potent repressor.

### The minimal enhancers of *D. sechellia A* and *DG2* retain activity

We next extended our analysis to the *A* and *DG2* enhancers to determine which evolutionary path underlies the reduced activity of these enhancers in *D. sechellia*. The *D. melanogaster A* (*melA*) enhancer drives expression in the cells that produce the thoracic tertiary trichomes and in lateral patches of the abdominal segments (Fig. 1L), and the *D. melanogaster DG2* (*melDG2*) enhancer drives expression in dorsal and lateral patches (Fig. 1H). Unlike *E6* and *Z1*.*3*, which have been dissected into small fragments that recapitulate the expression patterns of larger elements (*28, 36*), these enhancers reside in larger fragments that have not yet been subdivided into minimal elements. To uncover the genetic changes underlying their reduced activity in *D. sechellia* (Fig. 1I and M), we first sought to identify the minimal regions within the large *melA* and *melDG2* enhancers that sustain expression.

To identify the minimal region within the 5 kb *melA* enhancer that sustain expression, we leveraged published single-cell assay for transposase-accessible chromatin with sequencing (scATAC-seq) data from *D. melanogaster* embryos (*37*) and H3K27ac ChIP-seq data from *svb*-expressing cells (*38*). We used the scATAC-seq dataset to map regions of accessible chromatin in embryonic epidermal cells (Fig. 3). These analyses revealed that regions of open chromatin coincide with *svb* minimal enhancers: *Z1*.*3* (*28*), *E6* (*36*), and *7H* (*24*) (Fig. 3). These enhancer regions are also flanked by H3K27ac, a histone mark associated with active enhancers (Fig 3). For *A* and *DG2* we found a clear open-chromatin peak within the ∼5 kb of these elements. Guided by these data, we cloned a 1.2 kb fragment from the *A* enhancer, corresponding to the open-chromatin peak, and named it *melA1*.*2* (Fig 4A). This element drives higher levels of expression than of the full-length *melA* enhancer, with expanded activity in the abdominal tertiary trichome cells and broader expression in the lateral patches of the abdominal segments (Fig. 4J). Surprisingly, the orthologous sequence from *D. sechellia, secA1*.*2*, drove a comparable pattern and levels of expression to *melA1*.*2* (Fig. 4K-L). These findings suggest that *secA1*.*2* retains conserved activator binding sites, and that repressive elements acquired outside this minimal region are responsible for the reduced activity of the full-length *secA* enhancer.

**Fig. 3:**
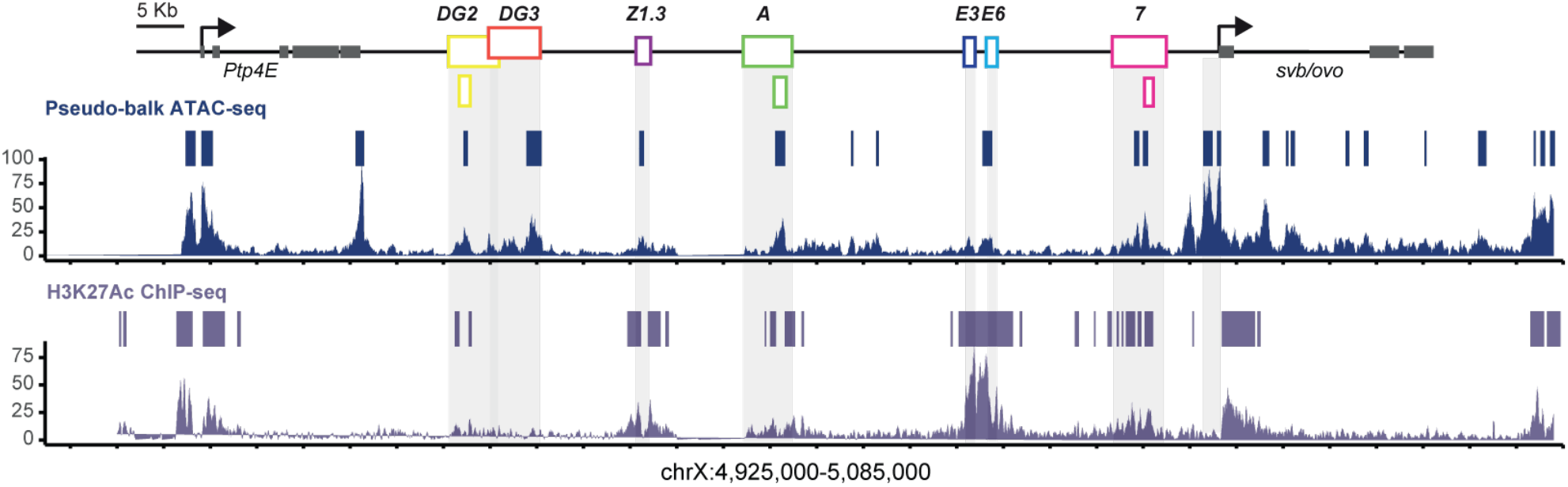
Chromatin landscape at the *svb* locus predicts minimal enhancers. Genome browser representations show pseudo-balk ATAC-seq profiles from the embryonic epidermis (top) and H3K27ac profiles from *svb*-expressing nuclei (bottom) across the *svb* genomic region. ATAC-seq and H3K27ac peaks are highlighted with blue and purple boxes, respectively, above each profile. Gray shading marks the positions of the *svb* enhancers, also shown in the schematic at the top. The locations of the *7H* enhancer (*24*)and the newly identified minimal enhancers are shown below this schematic.

**Fig. 4:**
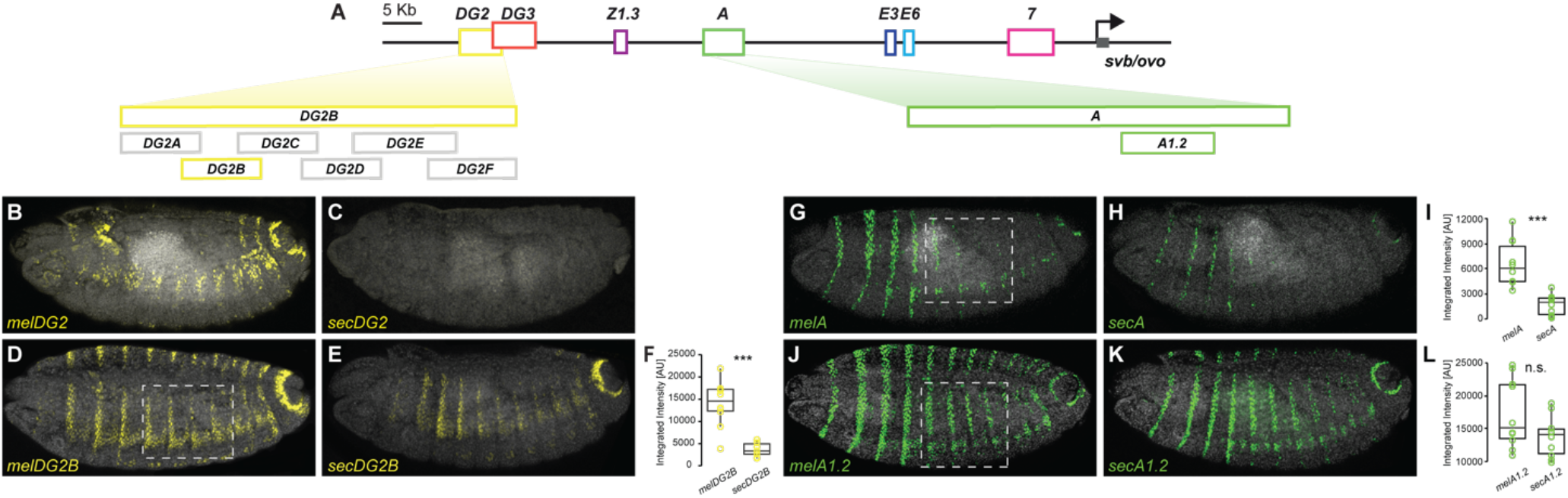
The minimal enhancer of the *D*.*sechellia DG2* evolved a reduced activity while the minimal enhancer of *D*.*sechellia A* encodes conserved activator binding sites. **(A)** Top: Structure of the *svb* locus, indicating the position of embryonic enhancers *svb* coding region is marked with gray arrow. Bottom: Schematics of the *DG2* and *A* enhancer fragments tested for epidermal enhancer activity in reporter gene assays. Gray fragment did not drive expression. **(B-E)** Expression of *melDG2* (B), *secDG2* (C), *melDG2B* (D), and *secDG2B* (E) reporter constructs in stage 15 embryos. **(F)** Box plots showing the integrated intensity of reporter activity in nuclei carrying the indicated constructs in the region outlined in B (n=10 embryos for each genotype). In the plots, each point represents an individual embryo. Asterisks denote significant difference between species, *** – P<0.001 (Student’s t test). **(G-L)** Expression of *melA* (G), *secA* (H), *melA1*.*2* (J), and *secA1*.*2* (K) reporter constructs in stage 15 embryos, juxtapose to box plots (I and L) showing the integrated intensity of reporter activity in nuclei carrying the indicated constructs as in the region outlined in G and J (n=10 embryos for each genotype). Each point in the plot represents an individual embryo. Asterisks denote significant difference between species, **** – P<0.0001, n.s. – not significant (Student’s t test).

We next used a systematic functional dissection to identify the minimal *melDG2* enhancer. We generated a series of overlapping fragments spanning the 5.2 kb *melDG2* region and identified a 1 kb fragment, named *melDG2B*, that retained expression similar to the full-length enhancer (Fig. 4D). All other fragments tested, did not drive any detectable expression in the embryo (data not shown). As expected, *melDG2B* corresponds to a region of open chromatin and is marked by H3K27ac in *D. melanogaster* embryonic epidermis (Fig. 3). We also examined expression driven by the orthologous *DG2B* sequences from *D. simulans* and *D. sechellia*. The *D. simulans* enhancer (*simDG2B*) drove expression similar to *melDG2B* (Fig. S3). Interestingly, the *D. sechellia* orthologous fragment, *secDG2B*, drove low levels of expression in a similar spatial pattern (Fig. 4E, 4F and S3). This observation suggests that, as with *E6* and *A, secDG2B* retains conserved activator inputs, while the full-length *secDG2* enhancer likely contains repressor binding sites that suppress its activity.

### The *D. sechellia A* enhancer acquired long-range repression upstream of the minimal enhancer

To identify the source of repression within the *D. sechellia A* enhancer, we tested larger enhancer fragments from both *D. melanogaster* and *D. sechellia* that included the minimal *A1*.*2* element and compared their activity (Fig. S4). All tested fragments, regardless of species origin, drove lower expression levels than the minimal *A1*.*2* constructs. This reduction may reflect conserved repressive inputs.

In all fragments that included sequences upstream of *A1*.*2*, the *D. sechellia* ortholog consistently drove significantly lower expression than the *D. melanogaster* ortholog. Moreover, the magnitude and significance of this difference increased with the length of the upstream region, with the most pronounced difference observed between the full-length *melA* and *secA* enhancers (Fig. S4). These findings suggest that repressive activity in *D. sechellia* is distributed across a broad genomic region upstream of *A1*.*2*, consistent with the presence of a long-range silencer element that evolved in the *secA* enhancer. Interestingly, the *secA* sequence contains an 833-bp insertion upstream of the *A3*.*6* fragment (Fig. S4), which may harbor some of this repressive activity.

### The *D. sechellia DG2* enhancer lost two activator binding sites

We next sought to identify the nucleotide changes responsible for the reduced function of the *D. sechellia DG2B* enhancer. First, we compared the *DG2B* sequence between *D. sechellia* and five related species, as was done for the *Z1*.*3* enhancer (Fig. S5). This analysis revealed 13 single-nucleotide substitutions, a three-nucleotide deletion and two single-nucleotide deletions unique to the *D. sechellia* sequence. To test the effect of these substitutions and deletions on enhancer activity, we introduced all of them into the *simDG2B* sequence and tested its function using quantitative reporter gene assay (Fig. 5). We chose to manipulate the *D. simulans* sequence because it is the species most closely related to *D. sechellia*. The resulting construct, designated *simDG2B_mutALL*, drove expression indistinguishable in pattern and levels from the *secDG2B* enhancer (Fig. 5E and F).

**Fig. 5:**
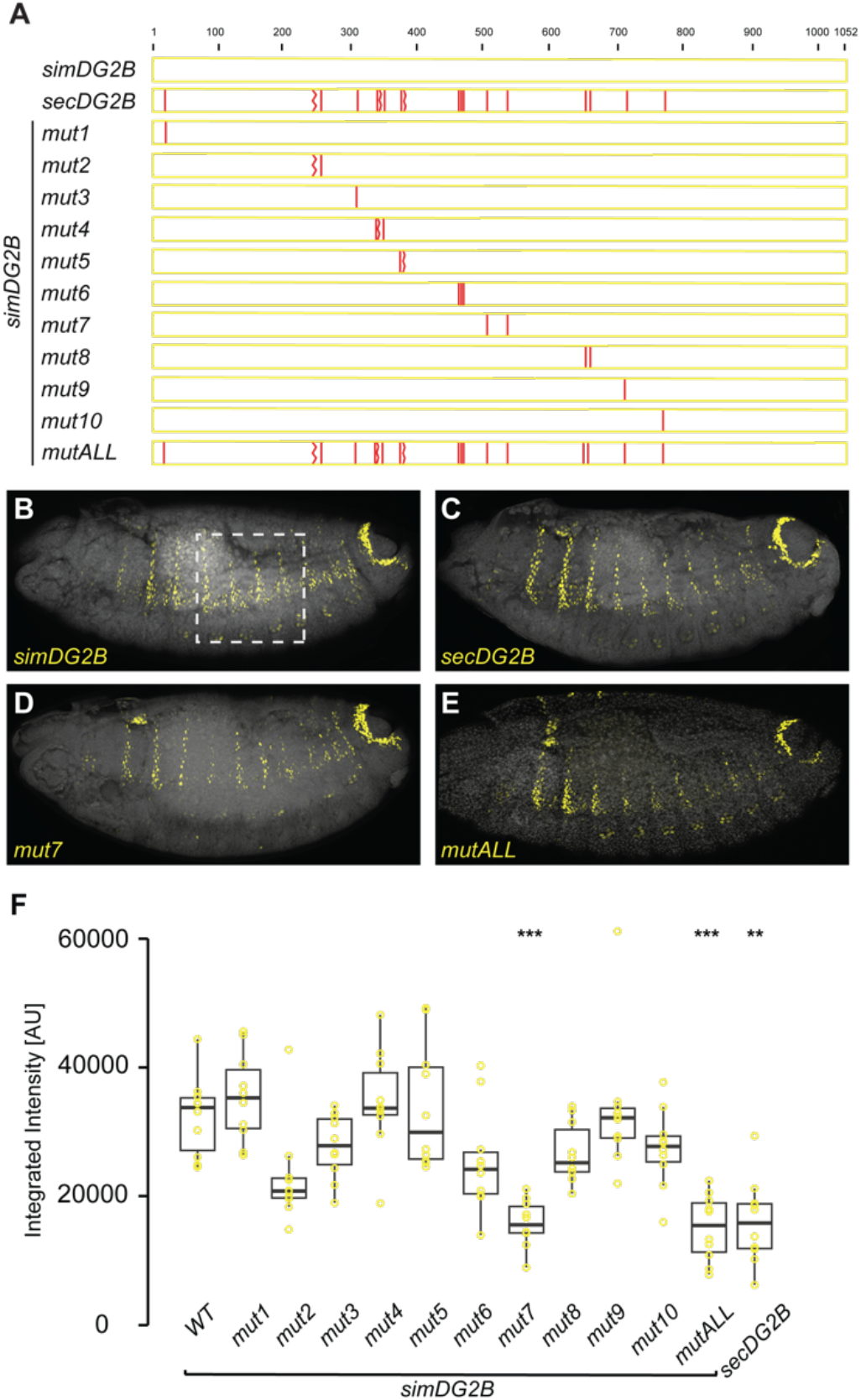
The *D. sechellia DG2* enhancer have lost two activator binding sites. **(A)** Schematic representation of the mutated *DG2B* enhancer reporter constructs. The positions of *D. sechellia*-specific substitutions are indicated with vertical red lines. **(B-D)** Expression of *simDG2B* (B) and *secDG2B* (C) reporter constructs and of *simDG2B* reporter constructs carrying the indicated *D. sechellia*-specific deletions and substitutions in stage 15 embryos. **(F)** Box plots showing the integrated intensity of reporter activity in nuclei carrying the indicated constructs in the region outlined in B (n=10 embryos for each genotype). In the plots, each point represents an individual embryo. Asterisks denote significant difference from *simDG2B* wild-type activities. (**) - P<0.01, (***) - P<0.001 (Kruskal–Wallis test with Hochberg-adjusted post hoc comparisons).

To determine which of the *D. sechellia*-specific nucleotide changes contributed to the reduced expression, we grouped them into ten clusters and introduced each cluster separately into the *simDG2B* sequence (Fig. 5A, S5 and S6). Strikingly, only one construct, named *simDG2B_mut7*, drove reduced expression levels, which are similar to *secDG2B* and *simDG2B_mutALL* (Figure 5C-F).

The *simDG2B_mut7* construct includes two *D. sechellia*-specific substitutions. We next asked which of these substitutions affect enhancer activity, and whether their effect results from the loss of activator binding sites or the gain of repressor binding sites. To investigate this, we conducted an adenine scan. We separately replaced the sequence surrounding each of the two sites with poly-A stretches in both the *simDG2B* and *secDG2B* enhancers (Fig. S7). The rationale for this experiment is that if a *D. sechellia* mutation caused the loss of an activator site, replacing the corresponding site in *simDG2B* with a poly-A stretch should reduce its activity without affecting *secDG2B*. Conversely, if the mutation introduced a repressor site, replacing it with a poly-A stretch should increase *secDG2B* activity while having no effect on *simDG2B*.

When we replaced each of the two candidate sites in *simDG2B* with poly-A stretches, both resulting constructs showed reduced enhancer activity (Fig. S7). However, only *simDG2B_polyA1* drove significantly lower expression levels compared to *simDG2B*, with levels comparable to those driven by *simDG2B_mut7*. In contrast, introducing the same poly-A substitutions into *secDG2B* did not significantly affect its activity (Fig. S7). These results suggest that the *D. sechellia*-specific substitutions primarily reduce *DG2B* activity by disrupting activator binding sites.

### Loss of activator sites unmasks conserved repression in the DG2 enhancer of *D. sechellia*

Our finding that the minimal enhancer *secDG2B* can drive residual expression, while the full *secDG2* enhancer does not drive any detectable expression, suggests that the *secDG2* enhancer contains repressive sequences outside of *secDG2B*. To identify the source of this repression, we tested larger fragments surrounding *DG2B* (Fig. 6). A fragment that included upstream sequences of *secDG2B*, named *secDG2AB*, drove similar expression to *secDG2B* (Fig. 6E and J). Interestingly, the orthologous sequence from *D. melanogaster, melDG2AB*, drove slightly lower expression compared to *melDG2B* (Fig. 6D and J), suggesting that this upstream region contains a repressor binding site specific to the *D. melanogaster* sequence.

**Fig. 6:**
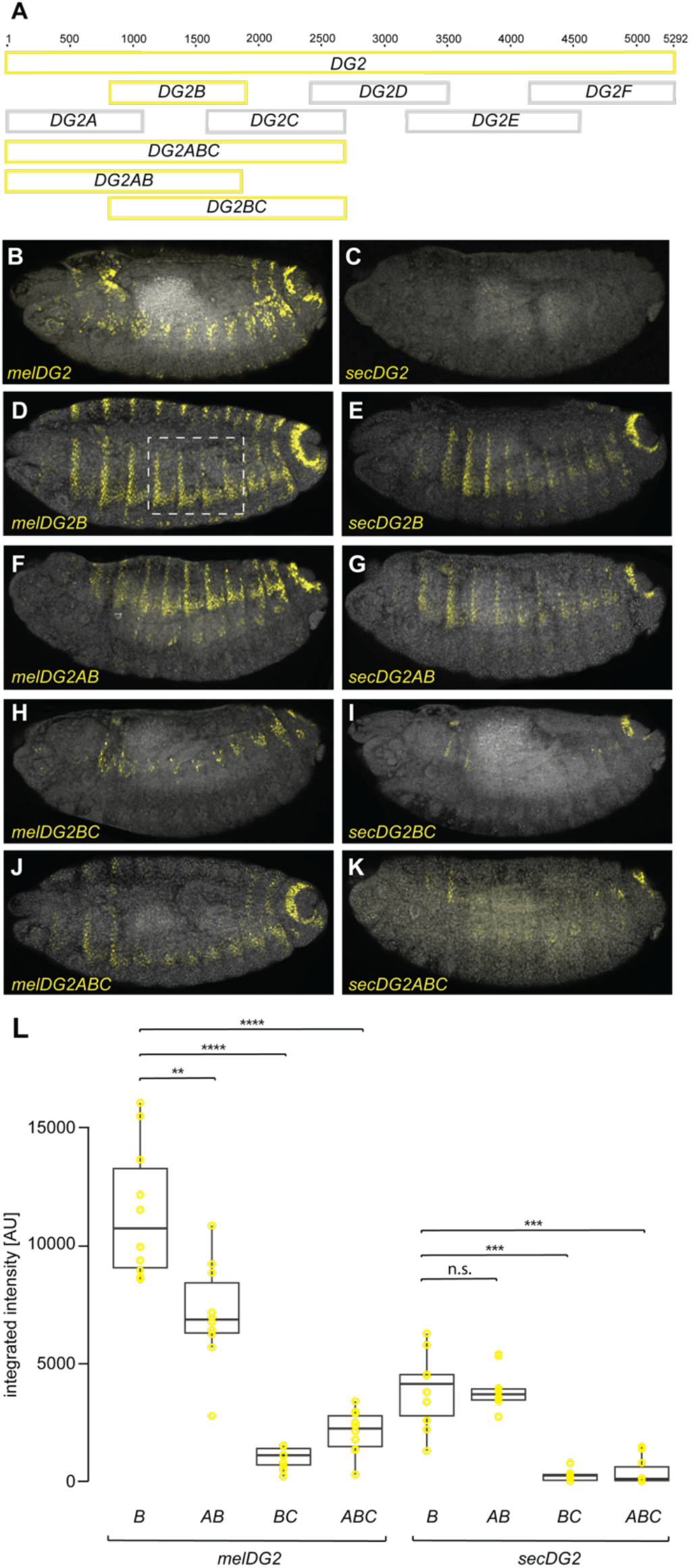
The *D. sechellia DG2* enhancer evolved through minimizing enhancer robustness to conserved repression. **(A)** Schematics of the *DG2* enhancer fragments tested for epidermal enhancer activity in reporter gene assays. Gray fragment did not drive expression. **(B-K)** Expression of the indicated *DG2* reporter constructs from *D. melanogaster* (B, D, F, H and J) and *D. sechellia* (C, E, G I and K)) in stage 15 embryos. **(L)** Box plots showing the integrated intensity of reporter activity in nuclei carrying the indicated constructs in the region outlined in D (n=10 embryos for each genotype). In the plot, each point represents an individual embryo. Asterisks denote significant difference from wild-type **– P<0.01, ***– P<0.001, **** – P<0.0001, n.s. – not significant (Kruskal–Wallis test with Hochberg-adjusted post hoc comparisons).

In contrast, the fragment *secDG2BC*, which included downstream sequences of *secDG2B*, showed markedly reduced activity, indicating the presence of repressive elements in this region (Fig. 6G and J). Surprisingly, the orthologous sequence from *D. melanogaster, melDG2BC*, also drove lower expression compared to *melDG2B*, suggesting that repression within the *DG2C* fragment is conserved across species (Fig. 6F and J). To account for the possibility that this reduction resulted from increased distance between the enhancer and the minimal promoter in the reporter construct, we inverted the enhancer orientation, so that the *DG2B* fragments remained adjacent to the promoter (Fig. S8). Interestingly, inverting *melDG2B* (*melDG2B*-*flipped*) and *secDG2B* (*secDG2B*-*flipped*) both led to reduced expression, suggesting that their activating elements are located near the end adjacent to the *DG2C* fragment, and that distance from the promoter does play a role (Fig. S8C, F, H and K). Nonetheless, adding the *DG2C* fragment upstream of *DG2B*-*flipped* from either species (*melDG2CB* and *secDG2CB*) led to a further reduction in expression (Fig. S8E, F, G and K). These results support the conclusion that the *DG2C* fragment contains conserved repressive sequences, although increased distance from the promoter may also contribute to the observed reduction in activity.

To further dissect the repressive activity within *secDG2C*, we divided the region into six ∼100 bp fragments and tested each one individually by placing it upstream of the *secDG2B-flipped* enhancer (Fig. S9). Four fragments (*secDG2C1, secDG2C2, secDG2C3*, and *secDG2C4*) reduced expression compared to *secDG2B-flipped* (Fig. S9Q), but only *secDG2C2* caused a significant decrease (Fig. S9K and Q). In contrast, none of the corresponding *melDG2C* fragments significantly lowered expression (Fig. S9P), suggesting that the repressive effect in *D. melanogaster* cannot be attributed to a short, isolated sequence.

These results indicate that in both species, the *DG2C* region contains repressive elements that reduce *DG2B* activity. We propose that the *secDG2* enhancer was inactivated through the loss of activator binding sites in *secDG2B*, making repressive sites in the *DG2C* region preponderant in the enhancer.

## Discussion

Genes controlled by redundant regulatory elements raise a central paradox: how can such robust expression systems be dismantled during evolution? Using the *Drosophila svb* gene as a model, we demonstrate that the evolutionary loss of quaternary trichomes in *D. sechellia* larvae occurred through the independent inactivation of four embryonic enhancers (Fig. 7). Each enhancer lost its function through a different molecular mechanism: single-nucleotide substitutions that removed activator binding sites and created a repressor binding site in *E6*, acquisition of long-range repression in *A*, loss of activator binding that unmasked conserved repression in *DG2*, and a large-effect deletion in the core of *Z1*.*3*. Notably, three of these mechanisms involved repression, highlighting repressive interactions as an underappreciated driver of regulatory change and suggesting that they may provide a particularly rapid path to phenotypic evolution.

**Fig. 7:**
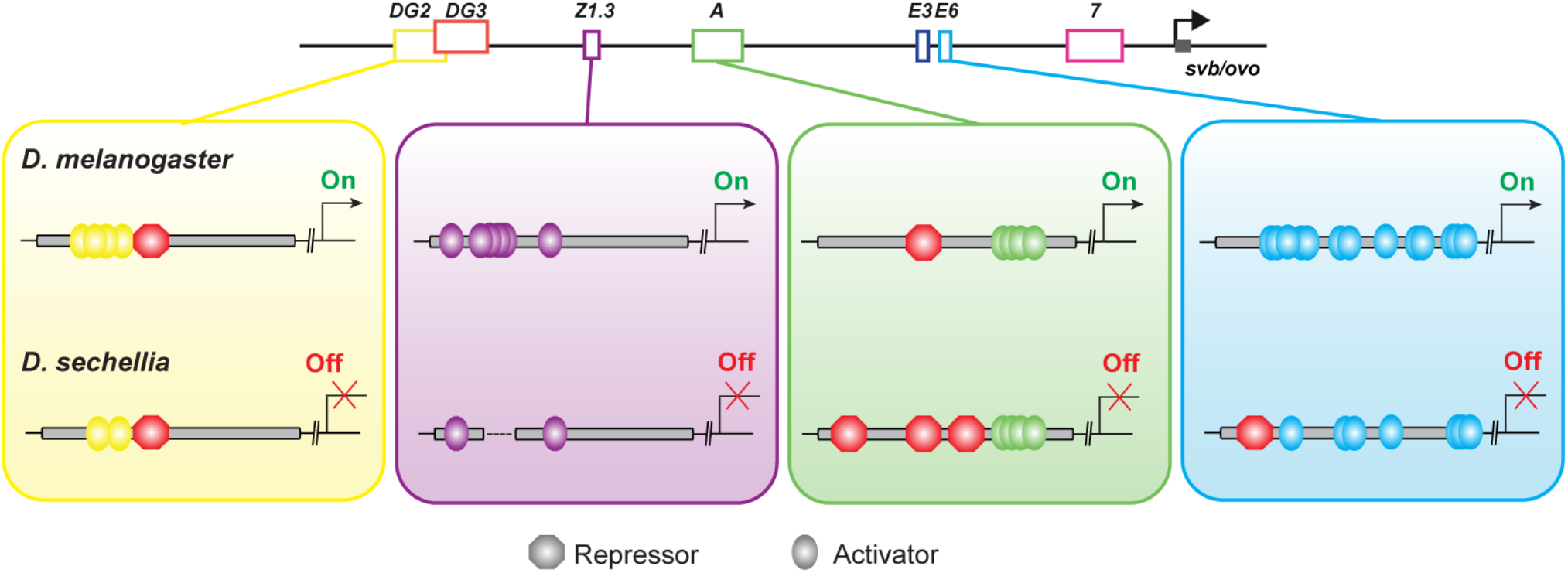
The *shavenbaby* enhancers evolved reduced functionality in *D. sechellia* through diverse mechanisms. Schematics illustrate predicted transcription factor occupancy at the *svb DG2* (yellow), *Z1*.*3* (purple), *A* (green), and *E6* (cyan) enhancers in *D. melanogaster* (top) and *D. sechellia* (bottom). Circles represent putative activator binding sites; octagons represent putative repressor binding sites.

Our work reveals that enhancer inactivation can occur through different molecular routes involving repression. We previously showed that the *svb E6* enhancer lost function in *D. sechellia* through the combined effect of multiple single-nucleotide substitutions that eliminated several Awh binding sites and created a novel binding site for the potent repressor Abrupt (*23*). This combination enabled complete inactivation of a previously robust enhancer. Here, we show that the *svb A* and the *DG2* enhancers were also inactivated through repression. The repeated evolution of repression across independent enhancers indicates that the naked cuticle of *D. sechellia* was shaped by natural selection rather than by genetic drift.

The *A* enhancer in *D. sechellia* was inactivated through the evolution of a long-range silencer that suppresses its activity. Silencers have previously been implicated in additional cases of morphological evolution in *Drosophila*. For example, gain and loss of silencer elements in the regulatory region of the *ebony* gene contributed to evolutionary changes in male abdominal pigmentation (*39, 40*). Similarly, the emergence of a silencer in the *yellow* gene regulatory region led to the secondary loss of wing spots in *D. melanogaster* (*11*). Our findings at the *svb* locus add a new example and suggest that modulation of silencer activities may be a common route for morphological change. Despite their importance, silencers remain an understudied class of regulatory elements, and their mechanisms of action are poorly understood (*41*). It remains unclear whether they act locally by interfering with enhancer activity or more broadly by inhibiting transcription at promoters. Addressing these questions will require genomic approaches such as ATAC-seq and chromatin conformation assays, which can provide insight into chromatin accessibility and long-range genomic interactions in an evolutionary context.

The *DG2* enhancer reveals a novel mode of inactivation: loss of resilience to conserved repression. While the minimal *secDG2B* fragment retains residual activity, the full-length *secDG2* is completely inactive. Dissection of adjacent sequences showed that the *DG2C* region carries repressive properties conserved between species. In *D. melanogaster*, this repression appears to fine-tune the expression domain, but in *D. sechellia*, the loss of activator input in *DG2B* renders the enhancer vulnerable to this pre-existing repression, resulting in complete silencing. To our knowledge, this is the first described case in which enhancer inactivation occurred through loss of activator binding that unmasked pre-existing repression.

The *Z1*.*3* enhancer is unique among the *svb* enhancers in that it was inactivated through a 120 bp deletion within its core sequence. Deletions of enhancer sequences have been documented in several cases of morphological evolution (*9, 42*–*44*), including the repeated loss of pelvic spines in freshwater stickleback populations (*9*). However, this type of mutation is thought to be less likely in enhancers that are pleiotropic, as deletions risk disrupting multiple regulatory functions. We previously showed that all embryonic *svb* enhancers can drive gene expression in multiple tissues and at different developmental stages (*28*). Unlike other *svb* enhancers, where the same transcription factor binding sites are reused across developmental contexts, the *Z1*.*3* enhancer exhibits a physical separation between the sequences required for embryonic and pupal expression (*28*). Strikingly, the 120 bp deletion in *D. sechellia* removes only the sequences necessary for embryonic activity, while leaving the adjacent pupal regulatory region intact. As a result, the pupal activity of *Z1*.*3* is preserved in *D. sechellia* (*28*). This organization of regulatory information thus permitted the selective loss of embryonic function through deletion, without compromising the enhancer’s roles in other developmental contexts.

While we have focused here on sequence-level changes within enhancers, the broader regulatory landscape of the *svb* locus also warrants attention. Given the extensive *cis*-regulatory inactivation observed in *D. sechellia*, an important open question is whether chromatin architecture has also evolved in *D. sechellia*. For example, how have the sequence changes in *D. sechellia* affected chromatin accessibility? On one hand, the acquisition of a silencer, as seen in *secA*, could increase chromatin accessibility, as silencers often exhibit open chromatin that enables repressor binding (*11*). On the other hand, potent repressors like Abrupt, which represses the activity of *secE6*, might function by promoting chromatin compaction, leading to reduced accessibility. These possibilities could be tested using comparative, cell-type-specific ATAC-seq, which may also help pinpoint the location and extent of repressive activity in the *D. sechellia A* enhancer.

Another timely question concerns the three-dimensional (3D) organization of the *svb* locus in *D. sechellia*. Do physical interactions between enhancers and the *svb* promoter persist in *D. sechellia* despite the loss of enhancer activity? Or has the loss of activator input and the acquisition of repression reshaped the 3D architecture of this locus? We have recently found that in *D. melanogaster*, the *svb* enhancers form a 3D hub that brings multiple enhancers into contact with the *svb* promoter (Naik et al., in preparation). Interestingly, this hub persists even in the absence of transcription factor binding to individual enhancers. This observation raises the possibility that the genetic changes affecting enhancer function in *D. sechellia* may not disrupt the higher-order chromatin structure of the locus. If so, this would suggest that 3D enhancer–promoter organization is robust to genetic perturbations of this scale. Testing this hypothesis will require cell-type specific chromosome conformation approaches that can directly compare the spatial organization of the *svb* locus between *D. melanogaster* and *D. sechellia* embryos.

Across metazoa, most morphological traits are governed by robust regulatory mechanisms that ensure phenotypic reproducibility, as many of their component parts are essential for survival. This study provides a detailed genetic account of how such robust systems, once thought to buffer against evolutionary change, can themselves serve as the substrate for morphological divergence. Our results show that evolution can exploit diverse mutational paths to silence gene expression: deleting enhancers, weakening their activity, or repressing them through existing or novel mechanisms. These findings underscore the flexibility of *cis*-regulatory evolution and reveal how robust developmental systems remain evolvable.

## Materials and Methods

### Transgenic Constructs and Fly Strains

Enhancer fragments were amplified by PCR from genomic DNA of the corresponding species (see Table S1 for details). For mutagenesis experiments, the *Z1*.*3* mutated fragments were synthesized by GenScript, and the *DG2B* mutated fragments were generated by site-directed mutagenesis using PCR, except for the *DG2BmutAll* fragment, which was synthesized by IDT. The *Poly-A DG2B* fragments were synthesized by Twist Bioscience. All enhancer fragments were cloned into the *placZattB* reporter plasmid using Gibson Assembly (see Table S2 for cloning details). Plasmids were integrated into the attP2 landing site by Rainbow Transgenic Flies.

### Embryo Staining and Image Analysis

Stage 15 embryos were collected, fixed and stained using standard protocols with mouse anti-βGal (1:500, Promega) and AlexaFluor 488 goat anti-mouse (1:500, Invitrogen) antibodies. Embryos carrying reporter constructs were imaged on a ZEISS LSM 900 Confocal Microscope. Image analysis and fluorescent intensity quantification was performed using Fiji (https://imagej.net/)(*45*). Briefly, confocal stacks were converted to maximum intensity projections, and background fluorescence was subtracted using a rolling-ball radius of 50 pixels. Nuclei in abdominal segments A2–A5 were segmented using the ‘Analyze Particles’ tool, and the mean fluorescence intensity of each nucleus was measured. Expression was quantified in two ways: at the single-nucleus level, by pooling all nuclear intensities across embryos and visualizing their distribution with violin plots; and at the embryo level, by summing the mean intensities of all segmented nuclei within each embryo to yield an Integrated Intensity value.

### Statistical analysis

All statistical analyses and data visualizations were performed using R (version 4.3.1, 2023) (Table S3). To assess the distribution of each dataset, we first tested for normality using the Shapiro-Wilk test, and for homogeneity of variances using Levene’s test. For normally distributed data with equal variances, we applied Student’s t-test when comparing two groups (e.g. the *A* enhancer fragments), and one-way ANOVA when comparing more than two groups. In cases where ANOVA indicated a significant difference (P < 0.05), we performed Tukey’s Honestly Significant Difference (HSD) post hoc test to identify which specific groups are different, while controlling for multiple comparisons. If variances were unequal, we used Welch’s t-test or Welch’s ANOVA with p-value adjustment Hochberg, as appropriate. For non-normally distributed data, we used the Kruskal–Wallis test for comparisons among more than two groups, followed by the Hochberg procedure to adjust P-values.

### Genomic data analysis

Single-cell ATAC-seq FASTQ files from *Drosophila* embryos (GEO: GSE101581)(*37*) were processed as pseudo-bulk profiles using a previously described pipeline (available at https://github.com/laiker96/fastq_to_bam) (*38*). Reads from epidermal cells (stages 14–15), as annotated by the Descartes portal (*37*), were pooled prior to peak calling. Peaks were identified using MACS2 (v2.2.9.1) (*46*) with parameters adjusted to account for Tn5 insertion bias.

Similarly, H3K27ac ChIP-seq data from sorted *E10::GFP*-positive epidermal nuclei (*38*) were processed using bbduk.sh for adapter trimming, aligned with BWA-MEM2, and peak calling was performed with MACS2 (v2.2.9.1) using default parameters to identify sharp enrichment peaks characteristic of H3K27ac enrichment at active enhancers and promoters. Input H3 DNA was used as a control for background subtraction.

## Acknowledgments

We thank Mark Rebeiz and Troy Shirangi for critical comments that improved the manuscript. We also thank members of the Preger-Ben Noon lab for helpful discussions. We thank OpenAI’s ChatGPT (GPT-5, 2025) for assistance with language editing of the manuscript.

## Author contributions

Conceptualization: EPBN, ASA, NF

Methodology: ASA, EPBN

Investigation: ASA, NS, EF, SN, SR

Supervision: EPBN

Writing—original draft: EPBN, ASA

Writing—review & editing: EPBN, ASA, NF

## Data and materials availability

All data are available in the main text or the supplementary materials.

## Supplementary Materials

Figs. S1-S9

Tables S1-S3

