## Supplementary Materials for "Distinct mechanisms decommission redundant enhancers to facilitate phenotypic evolution"

Areej Said-Ahmad *et al.*

**This PDF file includes:**

Figs. S1 to S9.

**Other Supplementary Materials for this manuscript include the following:**

Data S1 to S3.

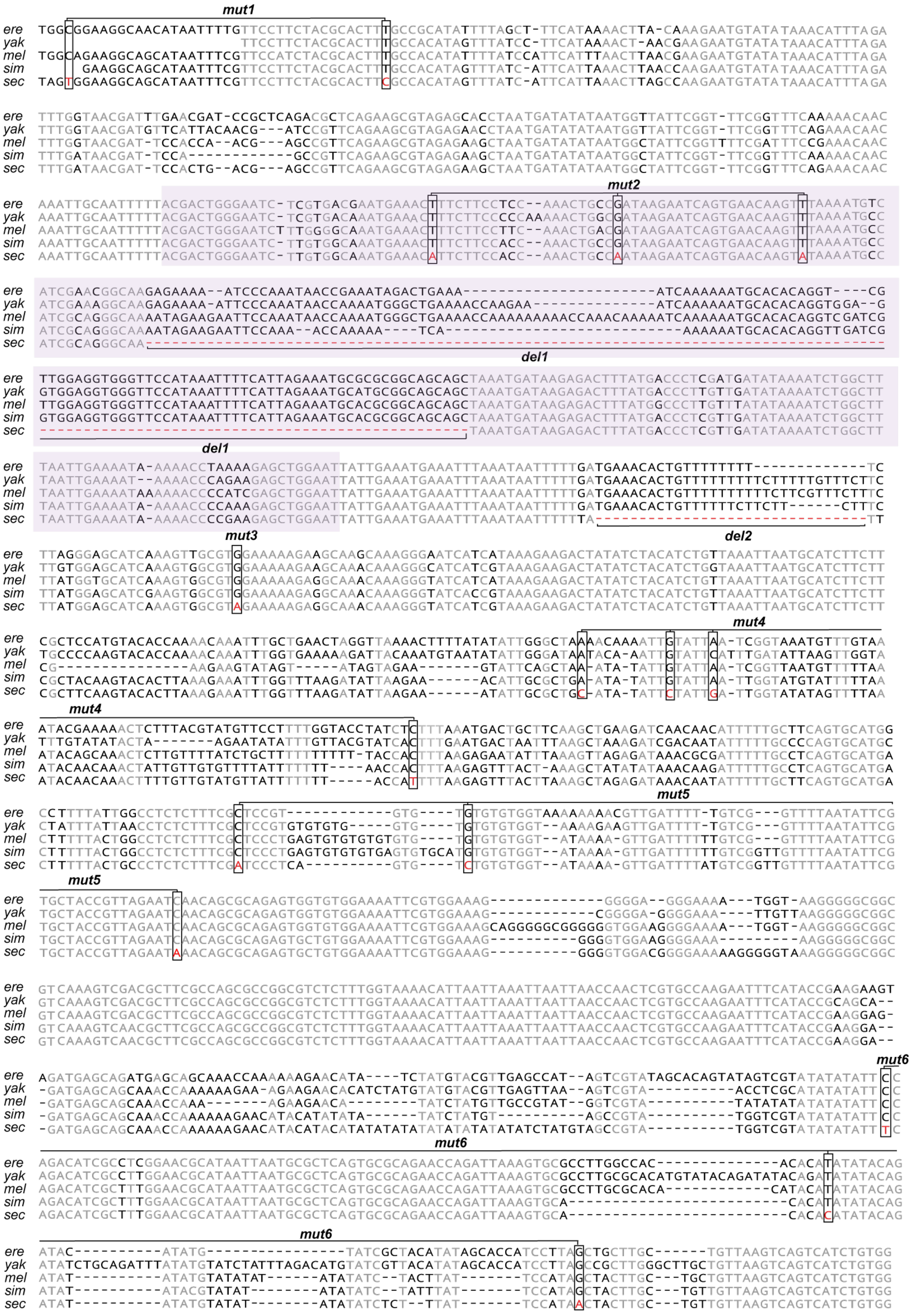
mut1
mut2
del1
mut3
del2
mut4
mut5
mut6
ere
yak
mel
sim
sec

**Fig. S1.**
Sequence alignment of the *Z1.3* region *D. erecta* (*ere*), *D. yakuba* (*yak*), *D. melanogaster* (*mel*), *D. simulans* (*sim*), and *D. sechellia* (*sec*). Conserved bases are shown in grey. Alignment positions with *D. sechellia*-specific substitutions are outlined with rectangles, and the substituted nucleotides are shown in red. The two *D. sechellia*-specific deletions are indicated, with deleted bases marked by red dashes. The location of the minimal enhancer *Z0.3* is highlighted in light purple. Clusters of mutations introduced into *melZ1.3* are indicated.

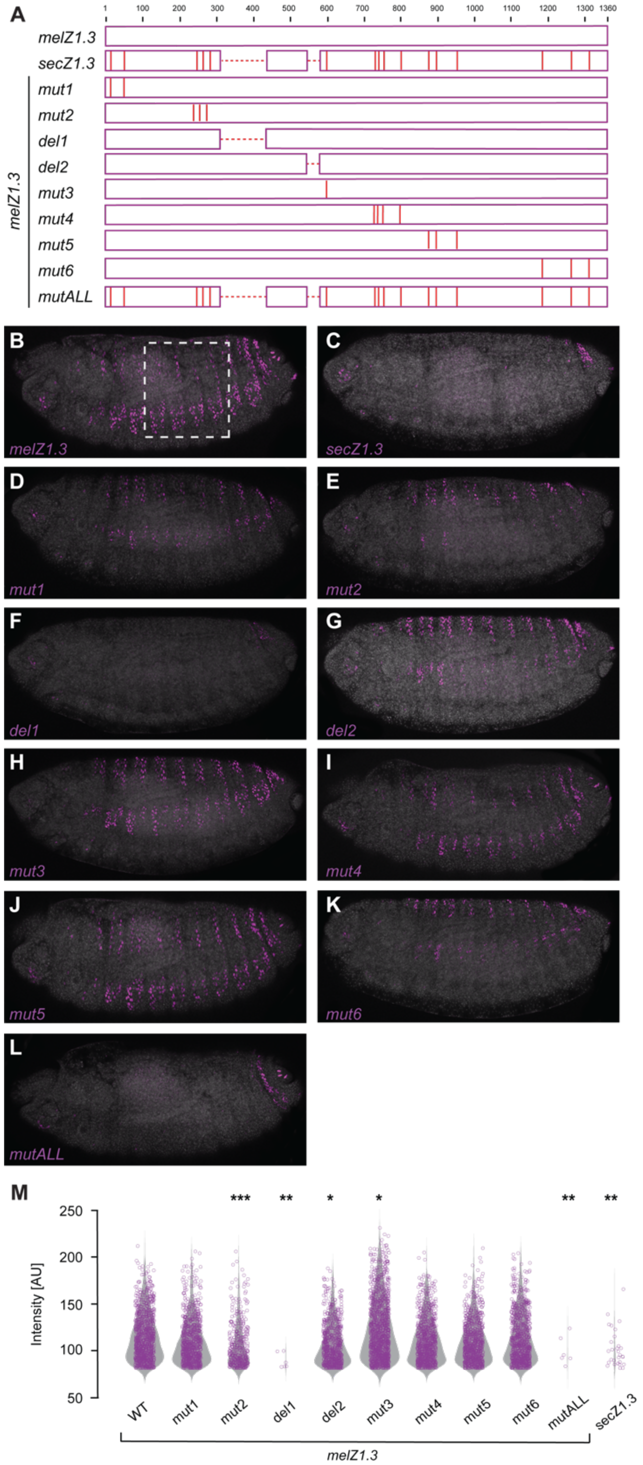


**Fig. S2.**
***One large effect deletion and three small effect nucleotide substitutions underly the loss of Z1.3 in D. sechellia.***
**(A)** Schematic representation of the mutated *Z1.3* enhancer reporter constructs. The positions of *D. sechellia*-specific substitutions and deletions are indicated with vertical and dashed red lines, respectively.

**(B-L)** Expression of *melZ1.3* (B) and *secZ1.3* (C) reporter constructs and of *melZ1.3* reporter constructs carrying the indicated *D. sechellia*-specific deletions and substitutions in stage 15 embryos.
**(M)** Violin plots showing the intensity of reporter activity in nuclei carrying the indicated constructs in the region outlined in B (n=10 embryos for each genotype). In violin plots, each point represents an individual nucleus. Asterisks denote significant difference from *melZ1.3* wild-type activities, * – $P<0.05$, **– $P<0.01$, ***– $P<0.001$ (Kruskal–Wallis test with Hochberg-adjusted post hoc comparisons).

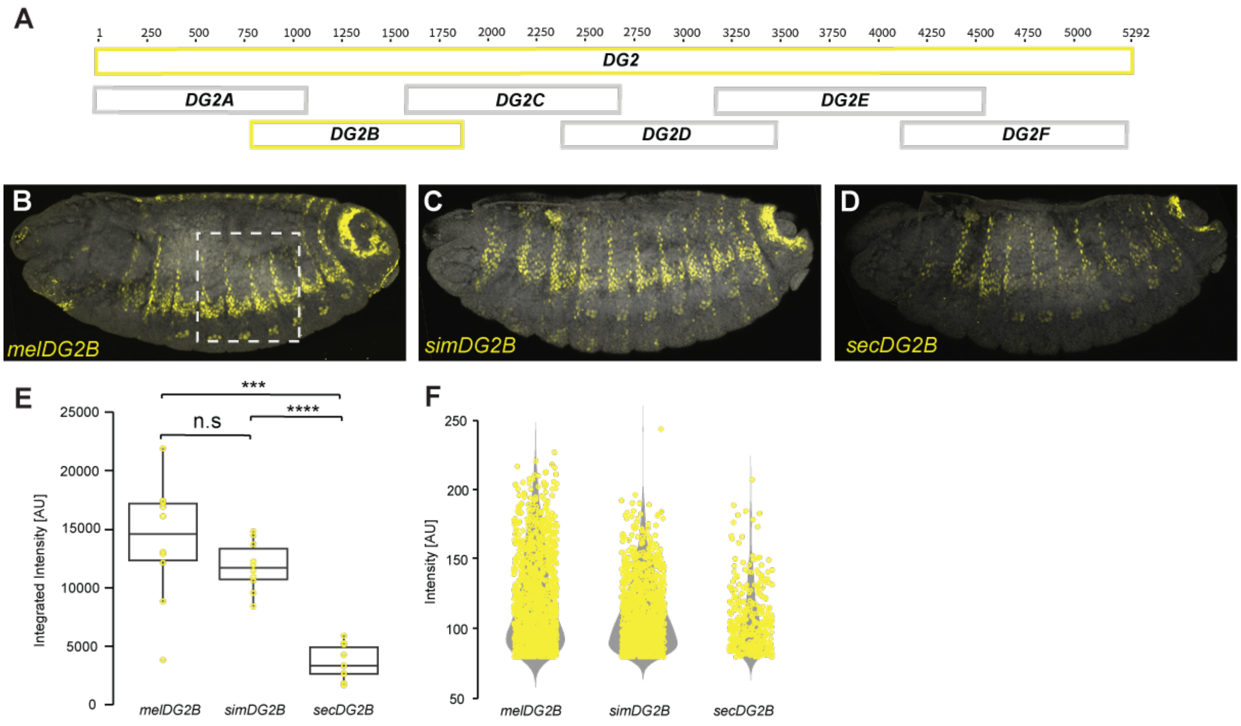


**Fig. S3.**
**The *D. sechellia DG2B* enhancer drives low levels of expression**
**(A)** Schematics of the *DG2* enhancer fragments tested for epidermal enhancer activity in reporter gene assay. Gray fragment did not drive expression.
**(B-D)** Expression of *melDG2B* (B), *simDG2B* (C), and *secDG2B* (D) reporter constructs in stage 15 embryos.
**(E-F)** Quantification of the intensity of reporter fluorescence in nuclei carrying the indicated constructs in the region outlined in B (n=10 embryos for each genotype). Each point in the box plot (E) represents an individual embryo while each point in the violin plots (F) represents an individual nucleus. Asterisks denote significant difference (****) - P<0.0001, (***) - P<0.001, n.s. – not significant (Welch's test).

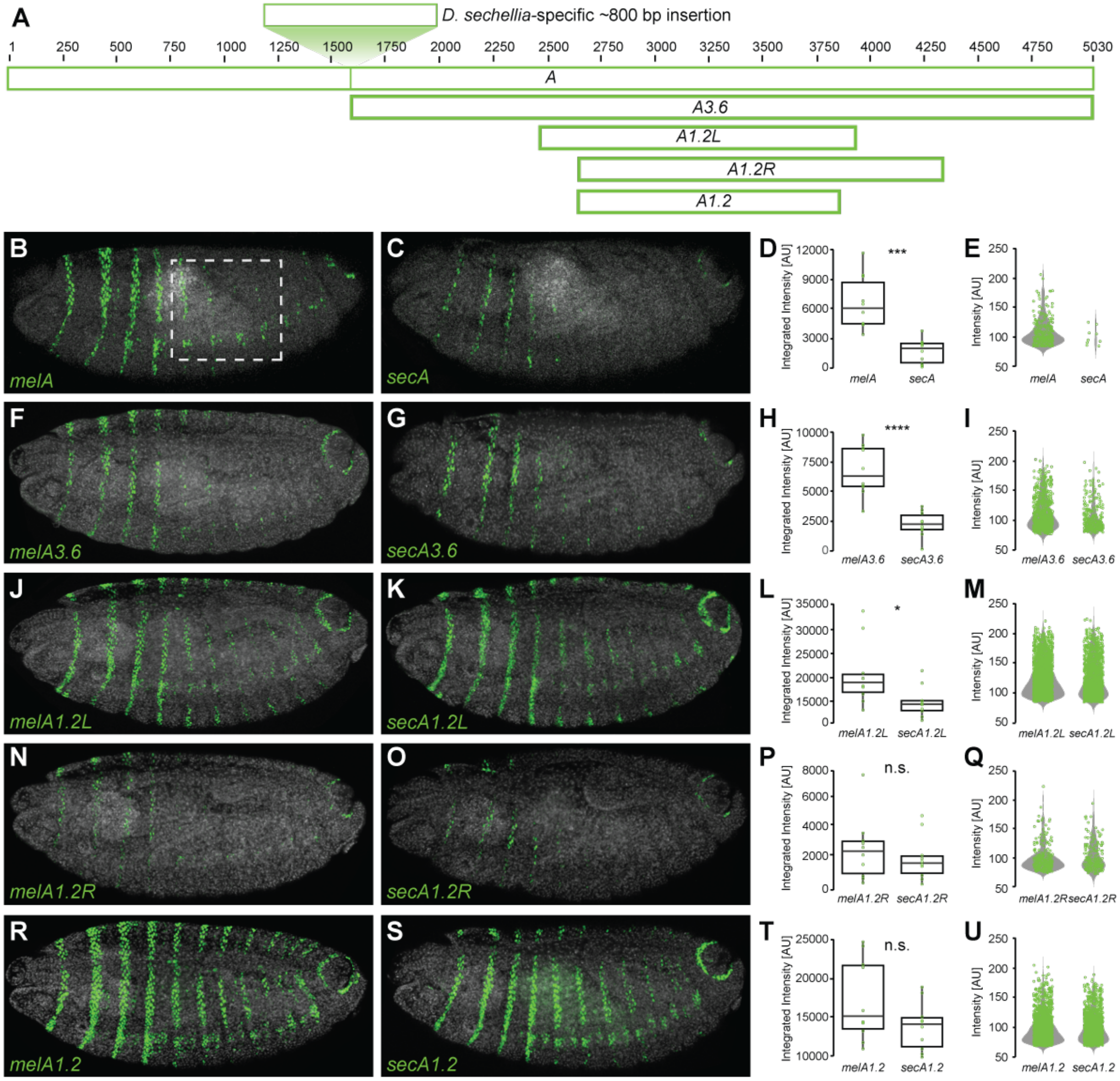


**Fig. S4.**

**The *A* enhancer in sechellia acquired repressive elements outside of the minimal enhancer *A1.2***

**(A)** Schematic representation of *A* fragments tested for epidermal enhancer activity in reporter gene assays. The location of a *D. sechellia*-specific ~800 bp insertion is indicated by a vertical line in the *A* fragment, and a representation of the inserted sequence is shown above it.

**(B-U)** Expression driven by the indicated *A* fragments, juxtaposed with box plots and violin plots showing the intensity of reporter activity in nuclei carrying the indicated constructs in the region outlined in B (n=10 embryos for each genotype). Each point in the box plot represents an individual embryo while each point in the violin plots represents an individual nucleus. Asterisks denote significant difference between species, *- P<0.05, ***– P<0.001, ****– P<0.0001, n.s. – not significant (Student's t test).

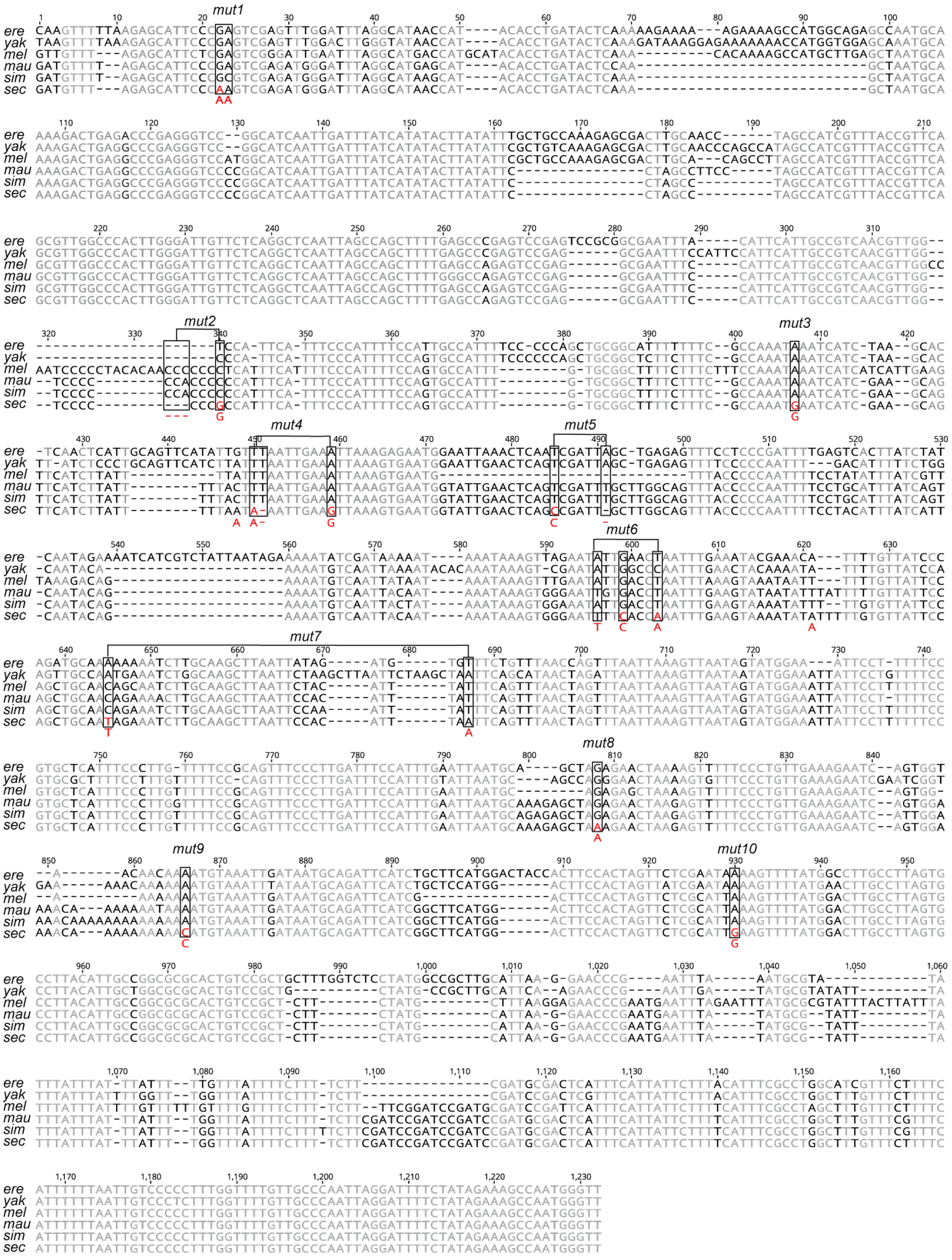
ere
yak
mel
mau
sim
sec
mut1
mut2
mut3
mut4
mut5
mut6
mut7
mut8
mut9
mut10

**Fig. S5.**

Sequence alignment of the *DG2B* region *D. erecta* (*ere*), *D. yakuba* (*yak*), *D. melanogaster* (*mel*), *D. mauritiana* (*mau*), *D. simulans* (*sim*), and *D. sechellia* (*sec*). Conserved bases are shown in grey. Alignment positions with *D. sechellia*-specific substitutions and deletions are outlined with rectangles, and the substituted nucleotides are shown in red. Clusters of mutations introduced into *simDG2B* are indicated, and the introduced mutations are shown below the sequence. In cases where a *D. simulans*-specific substitution was directly adjacent to a *D. sechellia*-specific substitution, the *simulans*-specific nucleotide was replaced with the *sechellia* base to preserve sequence context (e.g. *mut1*, *mut 4* and *mut6*).

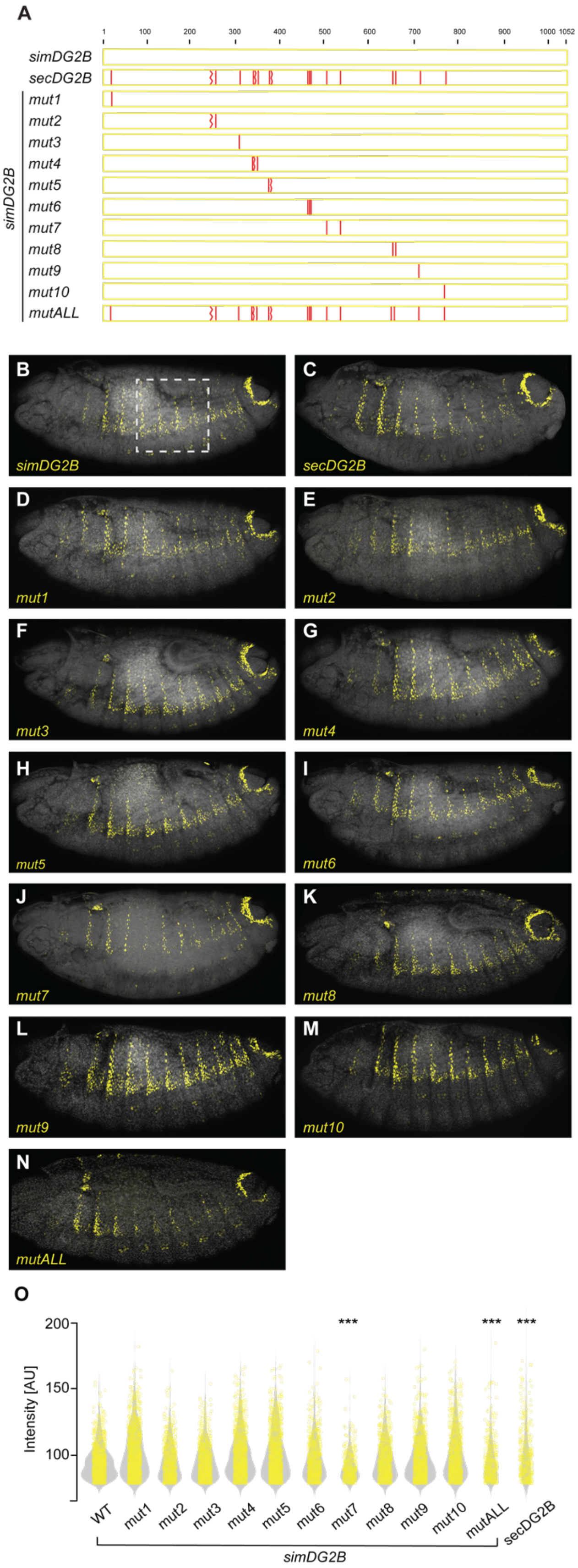
A
1
100
200
300
400
500
600
700
800
900
1000
1052
simDG2B
secDG2B
mut1
mut2
mut3
mut4
mut5
mut6
mut7
mut8
mut9
mut10
mutALL
simDG2B
B
simDG2B
C
secDG2B
D
mut1
E
mut2
F
mut3
G
mut4
H
mut5
I
mut6
J
mut7
K
mut8
L
mut9
M
mut10
N
mutALL
O
200
150
100
Intensity [AU]
***
***
***
WT
mut1
mut2
mut3
mut4
mut5
mut6
mut7
mut8
mut9
mut10
mutALL
secDG2B
simDG2B

**Fig. S6.**
**Two small effect nucleotide substitutions underly the reduced function of *DG2B* in *D. sechellia*.**
**(A)** Schematic representation of the mutated *DG2B* enhancer reporter constructs. The positions of *D. sechellia*-specific substitutions and deletions are indicated with vertical and zigzaged red lines, respectively.
**(B-N)** Expression of *simDG2B* (B) and *secDG2B* (C) reporter constructs and of *simDG2B* reporter constructs carrying the indicated *D. sechellia*-specific substitutions in stage 15 embryos.
**(O)** Violin plots showing the intensity of reporter activity in nuclei carrying the indicated constructs in the region outlined in B (n=10 embryos for each genotype). In violin plots, each point represents an individual nucleus. Asterisks denote significant difference from *simDG2B* wild-type activities. (**) - P<0.01, (***) - P<0.001 (Kruskal–Wallis test with Hochberg-adjusted post hoc comparisons).

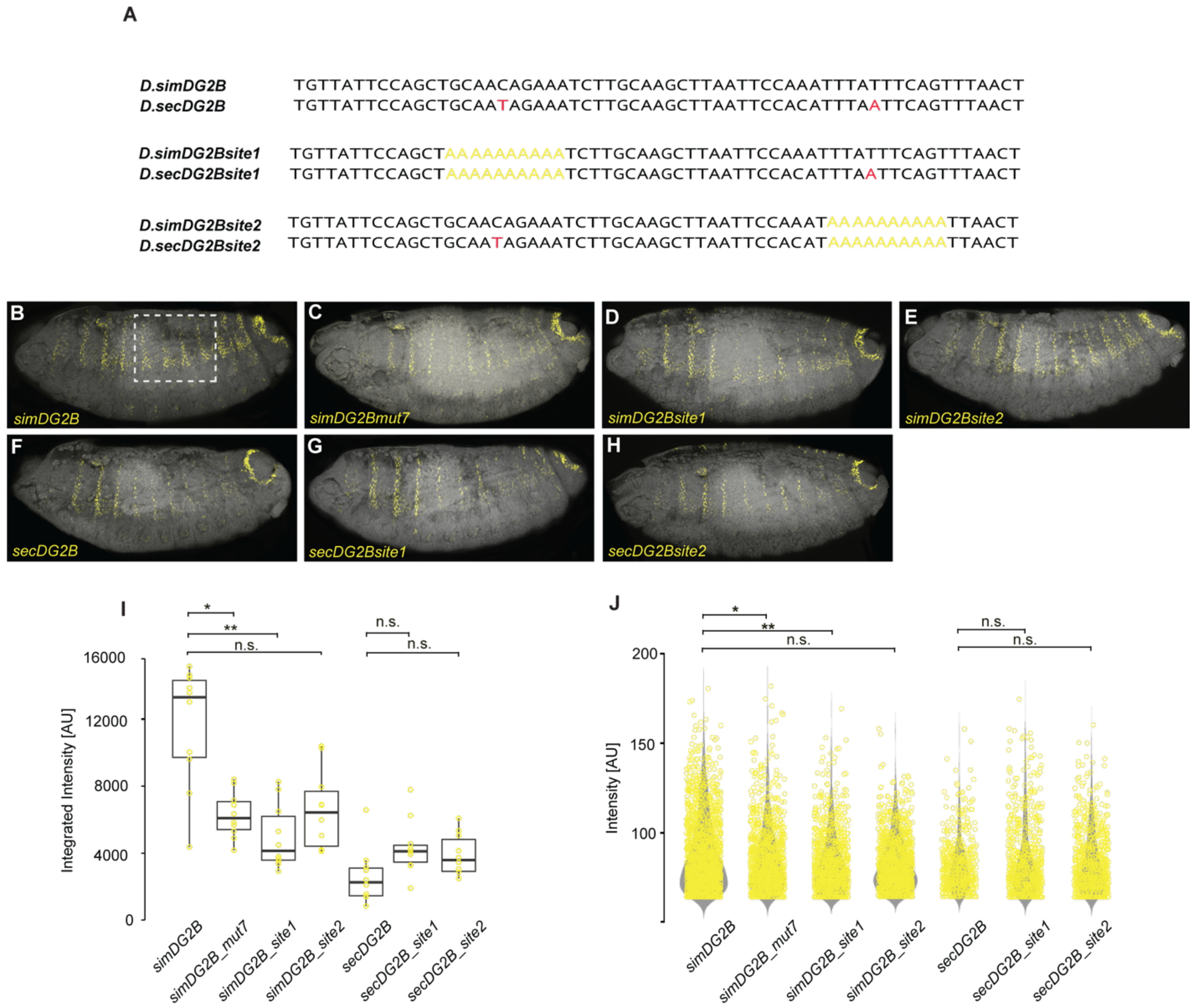


**Fig. S7.**

**The *D. sechellia DG2B* enhancer lost at least two activator binding sites**

**(A)** Top: Sequence alignment of the nucleotides surrounding the *D. sechellia*-specific substitutions in *mut7* from *D. simulans* and *D. sechellia*. The *D. sechellia*-specific nucleotides are marked in red within the *D. sechellia* sequence. Middle and bottom: Disruption of the sequences flanking each *D. sechellia*-specific substitution by replacing them with poly-adenine (poly-A) tracts, in the context of both *D. simulans* and *D. sechellia DG2B*.

**(B-H)** Expression driven by the indicated *DG2B* fragments in *D. melanogaster* stage 15 embryos.

**(I-J)** Quantification of the intensity of reporter fluorescence in nuclei carrying the indicated constructs in the region outlined in B (n=10 embryos for each genotype). Each point in the box plot (I) represents an individual embryo while each point in the violin plots (J) represents an individual nucleus. Asterisks denote significant difference from wild-type activities, * - P<0.05, **- P<0.01, n.s. – not significant (Kruskal–Wallis test with Hochberg-adjusted post hoc comparisons).

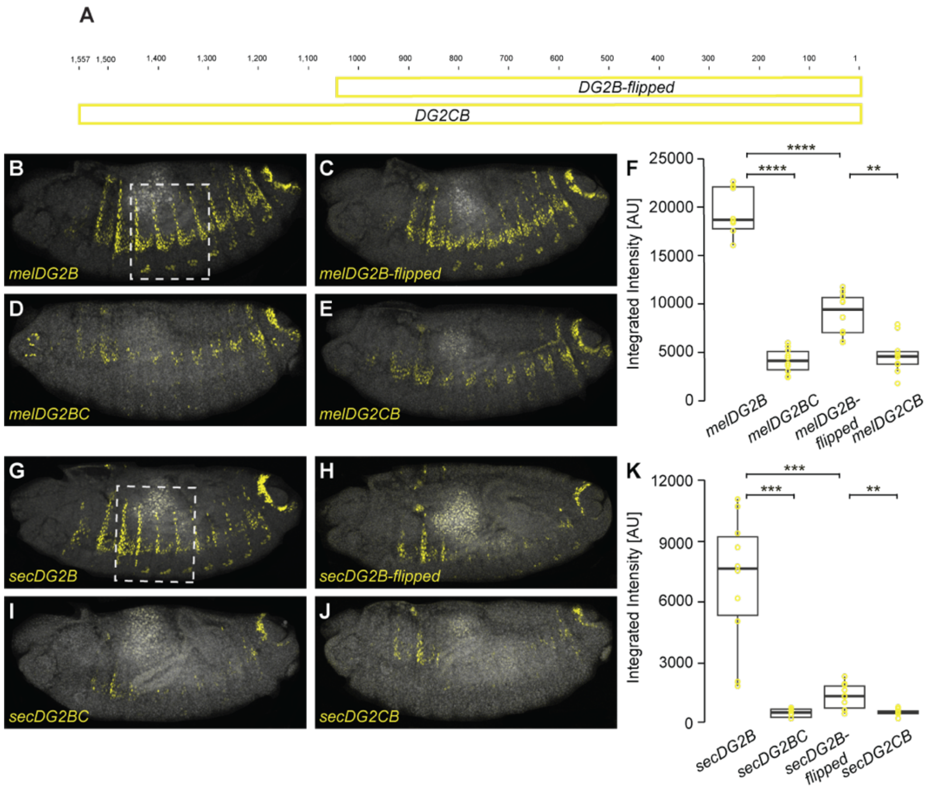


**Fig. S8.**
**The *DG2C* fragment contains conserved repression**
**(A)** Schematic representation of the flipped *DG2* enhancer reporter constructs.
**(B-E)** Expression driven by the indicated *D. melanogaster DG2* fragments in *D. melanogaster* stage 15 embryos.
**(F)** Quantification of the intensity of reporter fluorescence in nuclei carrying the indicated constructs in the region outlined in B (n=10 embryos for each genotype). Each point in the box plot represents an individual embryo. Asterisks denote significant difference between indicated samples, (**) - P<0.01, (***) - P<0.001, (****) - P<0.0001 (Kruskal-Wallis test and hochberg procedure for adjusting p-values).
**(G-J)** Expression driven by the indicated *D. sechellia DG2* fragments in *D. melanogaster* stage 15 embryos.
**(K)** Quantification of the intensity of reporter fluorescence in nuclei carrying the indicated in the region outlined in B, as in panel F.

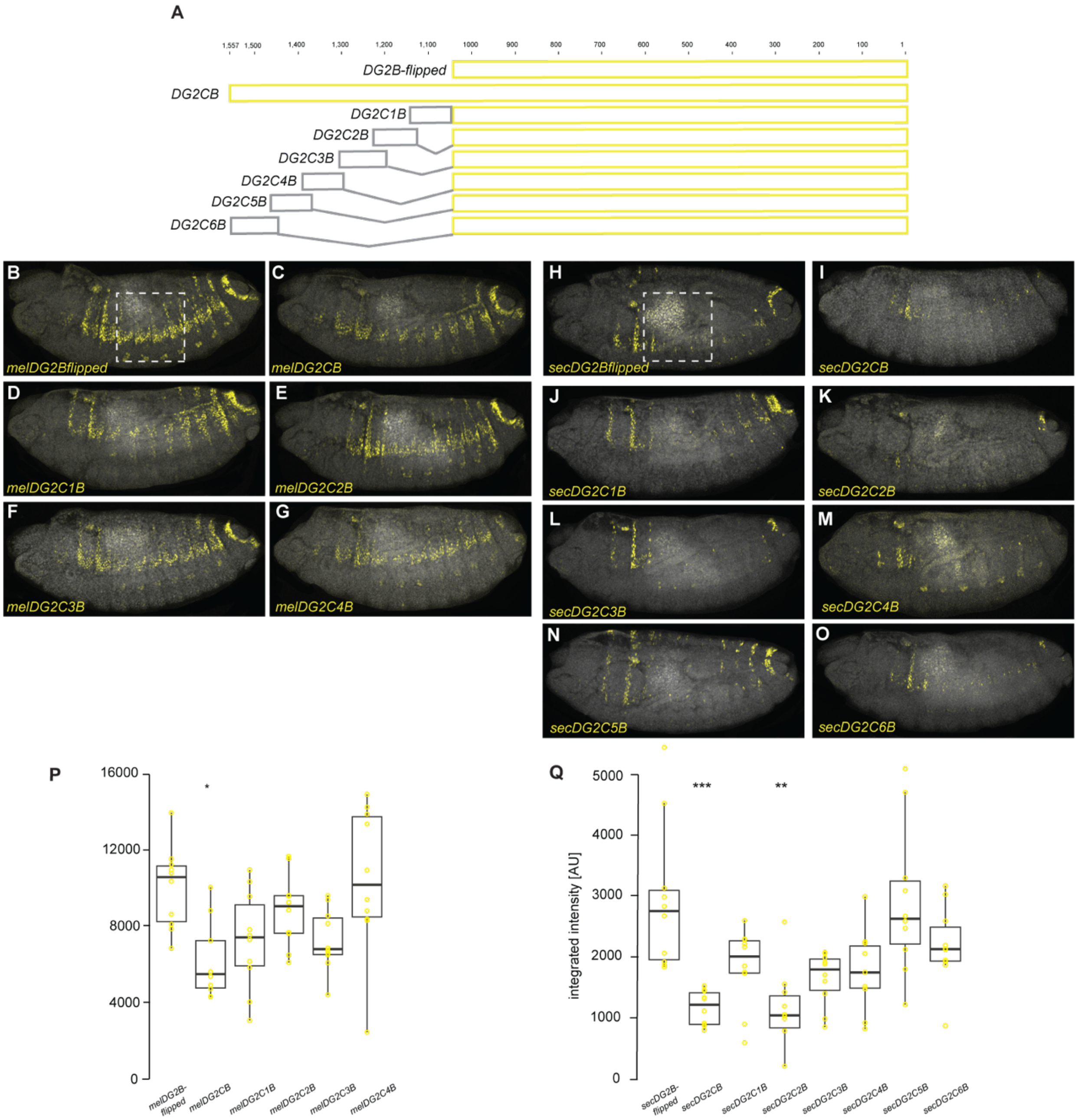


**Fig. S9.**
**The *DG2C* fragment contains conserved repression**
**(A)** Schematic representation of the chimeric *DG2B-flipped* reporter constructs. The *DG2B-flipped* fragment is shown in yellow. Attached *C* fragments, shown in gray, are positioned according to their relative location within the full *DG2C* sequence.
**(B-O)** Expression driven by the indicated *DG2B-flipped* reporter constructs in *D. melanogaster* stage 15 embryos.
**(P-Q)** Quantification of the intensity of reporter fluorescence in nuclei carrying the indicated constructs in the region outlined in B and H (n=10 embryos for each genotype). Each point in the box plot represents an individual embryo. Asterisks denote significant difference between

indicated samples, (*) - $P<0.05$, (**) - $P<0.01$, (***) - $P<0.001$(Kruskal-Wallis test and hochberg procedure for adjusting p-values).

**Data S1. (separate file)**

Table S1: List of primers used for cloning enhancer fragments and site-directed mutagenesis.

**Data S2. (separate file)**

Table S2: Sequences of enhancer fragments used in this study, with associated figures.

**Data S3. (separate file)**

Table S3: Summary of p-values from statistical comparisons between experimental groups.
